# A report on the detection and management of a finding of PSTVd (Potato Spindle Tuber Viroid) in potato breeding material in Northern Ireland, UK

**DOI:** 10.1101/728915

**Authors:** Richard O’Hanlon, Paul Watts, Rodney Martin, Gillian Young, Colin Fleming

## Abstract

Potato Spindle Tuber Viroid (PSTVd) is an infectious unencapsidated, small, circular, single-stranded RNA, that causes serious losses in infected potato plants. The pathogen has been detected in several European countries, including in England in the UK. In August 2016, the National Plant Protection Organisation of the Netherlands reported a finding of PSTVd in breeding material that originated in Northern Ireland, UK. A scheme of testing was carried out in Northern Ireland to identify the source of the infected breeding material. Trace-forward and trace-back tests identified 21 infected samples, of which 16 were from true seed samples, out of a total of 591 tested up to November 2016. The number of positive findings in true seed was further reduced to 4 after the samples were surface sterilized. Tests indicated that the infection probably entered the breeding station in Northern Ireland in the mid 1980’s, with limited spread in the collection via contaminated breeding equipment. The instance of spread in the field could not be ruled out. Eradication efforts included removal and destruction of infected field stocks and neighbouring stocks, destruction of stored museum material by deep burial, and destruction of other field material by ploughing and exposing tubers to frost. The risk of potato genitor material for spreading PSTVd is discussed. The situation of PSTVd in Northern Ireland, UK is Transient, actionable, under eradication.

## Introduction

Northern Ireland is recognised as a high grade seed potato producing area within the EU, and exports high grade seed potatoes (Pre-basic or Basic category seed of union grade PB, S, SE, or E Commission decision 2004/03/EC) to several countries. In total, the potato industry in Northern Ireland produced 119,200 tonnes of potatoes in 2018, with a value of over £20 million (DAERA 2019a). There are many plant health threats to potato in Northern Ireland, including the highly ranked threats from potato ring rot (causal agent Clavibacter michiganensis), root knot nematodes (*Meloidogyne fallax* and *Meloidogyne minor*) (DAERA 2019b). Protection of plant health is a devolved issue within the UK, with each of the four regions (England, Scotland, Wales and Northern Ireland) implementing and enforcing the EU plant health legislation in their own jurisdiction.

The pospiviroid Potato Spindle Tuber Viroid (PSTVd) is an infectious unencapsidated, small, circular, single-stranded RNA, capable of autonomous replication when inoculated into a host (see Jeffries 1998 for review). PSTVd is known to infect a wide range of crops in the wild, including potato, tomato, avocado (*Persea americana*), pepino, *Brugmansia* spp., *Solanum jasminoides*, pepper, and various other ornamentals (CABI 2017 webpage). It has a wide experimental host range, infecting 94 plant species in 31 families (Jeffries 1998). It is highly damaging to potato crops (EFSA 2011), and is regulated as an annex IAI quarantine pathogen in the EU, with legislation in place to prevent its introduction into and spread within the EU (2000/29/EC). Despite this legislation, PSTVd has been detected in Austria (first record 2008), Belgium (2006), Croatia (2009), Czech republic (2009), Germany (1980’s), Greece (2009), Hungary (2013), Italy (2007), Netherlands (2001), Poland (2016), Slovenia (2006), Spain (unknown), and England (2003) (EPPO GD 2017).

In August 2016 the NPPO of the Netherlands notified the NPPO for the UK (Department of the Environment, Food and Rural Affairs, DEFRA) that PSTVd had been detected in potato genitor material sent from Northern Ireland to the Netherlands (EPPO RS 09/2016). These genitor lines were not part of the commercial breeding programme in Northern Ireland and were only used as salt-tolerant parental breeding material. This report from the NPPO of Netherlands led to an extensive scheme of testing of related lines of potato in the Northern Ireland potato breeding programme, managed by the Agri-Food and Biosciences Institute (AFBI) potato research centre in Loughgall, Northern Ireland (hereafter AFBI Loughgall) and any commercial premises which had links with AFBI. The details of the detection, and subsequent testing programme are documented here.

## Methods

### Incident management team establishment

Following informal notification from the Dutch recipient of the breeding material of the detection of PSTVd to AFBI Loughgall, an Incident Management Team (IMT) was established to coordinate a response to the detection of the quarantine organism. The aim of the IMT was to identify the source of the PSTVd, eradicate PSTVd from Northern Ireland and facilitate any similar actions in the devolved administrations of the UK and any other EU member states. The IMT included:

- members of the Northern Ireland Department of Agriculture, Environment, and Rural Affairs (DAERA) with responsibility for plant health regulation
- the programme manager of the AFBI Loughgall potato breeding programme
- members of the AFBI Plant Health testing laboratory at Newforge lane, Belfast (hereafter AFBI Newforge).

The team met 12 times between September 2016 and May 2017 to review the situation and plan actions. The NPPO of the UK (DEFRA) was kept informed of all developments, as well as other NPPO’s when the need arose.

### Sample selection and testing

The IMT carried out a series of trace-back and trace-forward searches on all potato breeding material in the Northern Irish potato breeding programme, starting at the genitor material found to be positive by the Netherlands NPPO. Samples of tubers and true seed were identified and sent to AFBI Newforge for testing. Testing followed the protocol outlined in the International Standards for Phytosanitary Measures (ISPM 27 Diagnostic Protocols). Potatoes were first washed in tap water and dried at room temperature. A single core (1 cm diameter) was removed from the heel end of each tuber and bulked up to 200 cores per sample following the procedure used for Brown Rot/Ring Rot testing (EU directive 2006/63/EC). True seed was also tested, initially without surface sterilisation before testing. However, following detection of low levels of PSTVd in some true seed samples early in the testing scheme, the protocol was amended to include surface sterilisation. This involved incubating seeds for 5 minutes in 5ml ethanol (70%), washing in 5% sodium hypochlorite for 10 minutes followed by 2 × 5 minute washes in 15 ml sterile deionised water. The samples were tested using the real time q-PCR protocol of (Boonham *et al.*, 2004). DNA of one of the earliest detected infected samples (D1749, a cross between DT028 × Cara) was sequenced using 3H1 forward and 2H1 reverse primers (Shamoul et al. 1999) by a commercial sequencing service at Queens University Belfast Genomics Core Technology Unit using an Applied Biosystems 3730 (Thermo Fisher Scientific). The obtained sequences were compared using Basic Local Alignment Search Tool (BLAST) analysis with sequences available in GenBank (Altschul et al. 1990). The sequence was deposited in GenBank under the accession number MN231648.

## Results

Between the first detection of the organism on 31/08/16, and 10/11/16, a total of 591 samples were tested for the presence of PSTV. These included 355 samples from the field crop in AFBI, 45 from the glasshouse production facility at AFBI and 144 true seed samples from AFBI (Table 1). Testing was also carried out on 25 approved stocks in cases where the grower had recently received material from AFBI. Overall just 21 samples were positive for PSTVd, with 16 samples of true seed and 5 samples from the AFBI field crop testing positive (Table 1). Following surface sterilization of the true seed and re-testing, the number of positives from this source was reduced to 4.

**Table 1.** Details of sample numbers and PSTVd positive samples from the true seed before and after surface sterilization.

| Year seed was produced/imported | Number of samples tested | PSTVd positive samples | Positive samples after surface sterilization |
| --- | --- | --- | --- |
| 1984 | 14 | 0 | 0 |
| 1985 | 19 | 3 | 1 |
| 1991 | 12 | 1 | 1 |
| 1992 | 52 | 11 | 2 |
| 1994 | 1 | 0 | 0 |
| 1995 | 1 | 0 | 0 |
| 1996 | 2 | 0 | 0 |
| 1999 | 2 | 0 | 0 |
| 2001 | 5 | 0 | 0 |
| 2003 | 4 | 0 | 0 |
| 2004 | 5 | 0 | 0 |
| 2006 | 3 | 0 | 0 |
| 2007 | 11 | 0 | 0 |
| 2008 | 2 | 0 | 0 |
| 2009 | 1 | 0 | 0 |
| 2014 | 10 | 1 | 0 |
| Total | 144 | 16 | 4 |

### Trace-back testing

Following notification from the Netherlands that J3911/5 was positive for PSTVd, samples of the same cross, and other crosses that were in the crossing house in AFBI Loughgall at the time of production of the infected seed cross were tested. Of the four positive true seed samples after surface sterilization, two were the progeny of a salt-tolerant true seed sample DT028. The tuber sample of DT028 was tested twice in 2016 but found to be negative for PSTVd. However, this sample had been stored beside another seed sample (DT02) in the museum collection in AFBI since 1984. Although DT02 was not available for testing, the progeny (DT02 × Maris piper) of a cross carried out in 1985 using DT02 resulted in a PSTVd positive true seed sample after surface sterilization (Table 2). The final true seed sample (Dundrum × L2678/96), that was positive after surface sterilization, was potentially infected via one of its parents (L2678/96) which was used in the crossing house at the same time as a putatively infected source (DT028) in 1991.

**Table 2.** Timeline of events leading to the spread of PSTVd infection in the AFBI Loughgall potato breeding programme

| Date | Event | Details of samples tested | Result | Notes |
| --- | --- | --- | --- | --- |
| Unknown | DT02 received by Institute 1 | Not tested by PCR as material not available | N/a | N/a |
| 1981 | DT02 received by Institute 2 | Not tested by PCR as material not available | N/a | DT02 would have been tested for PSTV as part of the testing scheme at the Institute |
| 1983 | DT02 received by Institute 3 | Not tested by PCR as material not available | N/a | N/a |
| 1984 | Received by AFBI, Loughall | Not tested by PCR as material not available | N/a | N/a |
| 1985 | Used in crossing at Loughall. | DT02 X Maris Piper true seed. | PSTVd positive (2016) | Infection of true seed by infected pollen |
| 1985-1992 | DT02 maintained beside DT028 in | Not tested by PCR as material not available | N/a | Transmission of PSTV from DT02 to |
|  | museum<br>collection at<br>Loughgall. |  |  | DTO28<br>occurred in the<br>museum. |
| 1991 | DTO28 X<br>L2678/96 | True Seed | PSTVd positive<br>(2016) | Infection of true<br>seed by infected<br>pollen |
| 1992 | DTO28 X Cara | True Seed | PSTVd positive<br>(2016) | Infection of true<br>seed by infected<br>pollen |
| 2012 | DTO28 X Cara<br>true seed<br>germinated. | Not tested by<br>PCR as material<br>not available | N/a | N/a |
| 2013 - 2014 | DTO28 X Cara<br>crosses in<br>glasshouse &<br>polytunnel | Not tested by<br>PCR as material<br>not available | N/a | N/a |
| 2014 - 2016 | DTO28 X Cara<br>crosses<br>maintained in<br>field at<br>Loughgall | 2 salt tolerant<br>DTO28<br>x Cara crosses | PSTVd positive<br>(2016) | N/a |

In order to identify if any transmission had occurred in the crossing house during production of the infected line (J3911/5), and also to pinpoint which of the parents (DT028 or Cara) were the source of the infection, a scheme of testing was carried out (Table 1 and 2). Tests of true seed from crosses carried out in 1991 between DT028 × Cara detected PSTVd in one batch of seed before and after surface sterilisation. Another sibling cross from these same parents, J3911/6, was not available for testing but had been used to produce further crosses in 2014. All resulting crosses using J3911/6 (n=3), and any other crosses that were in close proximity to this cross in the crossing house in 2014 (n=4) were tested and found to be negative. In order to find the source of the infection in the J3911 siblings, other available crosses using DT028 and Cara were tested. A total of 23 crosses resulting from Cara were tested, of which 4 were positive before surface sterilization, and 0 positive after surface sterilization. All of the positive Cara crosses were produced in the crossing house in 1992. A total of 3 crosses that used DT028 were tested, of which 2 (DT028 × Cara; DT028×L2678/96) were positive. These infected crosses had been produced in the crossing house in 1991 and 1992. Tests of other true seed from crosses carried out in 1992 between DT028 × L2678/96, and between Dundrum × L2678/96 detected the pathogen before and after surface sterilization of the seed (Table 2). The museum collection at AFBI Loughgall was destroyed in February 2017 by deep burial.

### Trace forward testing

As part of the AFBI Loughgall potato breeding activity, field trials were carried out with varieties to assess their characteristics in the field. The genitor material originally notified by the Netherlands NPPO (J3911/5) was planted in field trials in 2016. The field crop covered an area of 3.4Ha of which approximately 60% was planted with potatoes. The field was divided into two main plantings corresponding to field slopes and to an original amalgamation of two smaller fields. The field was further sub-divided into 17 blocks with un-planted pathways to allow vehicle movements with minimal contact with foliage. The area containing the infected individual (J3911/5) notified by the Dutch NPPO was found in Block 2 (Figure 1), which was planted with advanced breeding lines and varieties in the process of registration. Within this block extensive testing detected a further 4 positive breeding lines based on tuber sampling. A sibling line to J3911/5, namely J3911/2 was also positive for PSTVd. Further positives were identified in three other breeding lines that were also located in block 2. All of the positive samples from the field were found in block 2. The infected lines and other stocks in close proximity to these lines were harvested, removed from field and destroyed by autoclaving at 121°C for 15 minutes and/or by deep burial. The remainder of the lines in Block 2 and adjacent blocks 1, 3 and 4 were destroyed by bringing the tubers to the surface using a rotavator to expose them to frost and rain. The field was sown with forage grass in 2017. Monitoring of the field in 2017 identified 6 volunteer potato plants, which tested negative for PSTVd and were treated with herbicide. The field was inspected again in July and November 2018 when no volunteer potato plants were found. Continued monitored of the field for volunteers is planned, and the field will not be used for potato production in the conceivable future.

**Figure 1.**
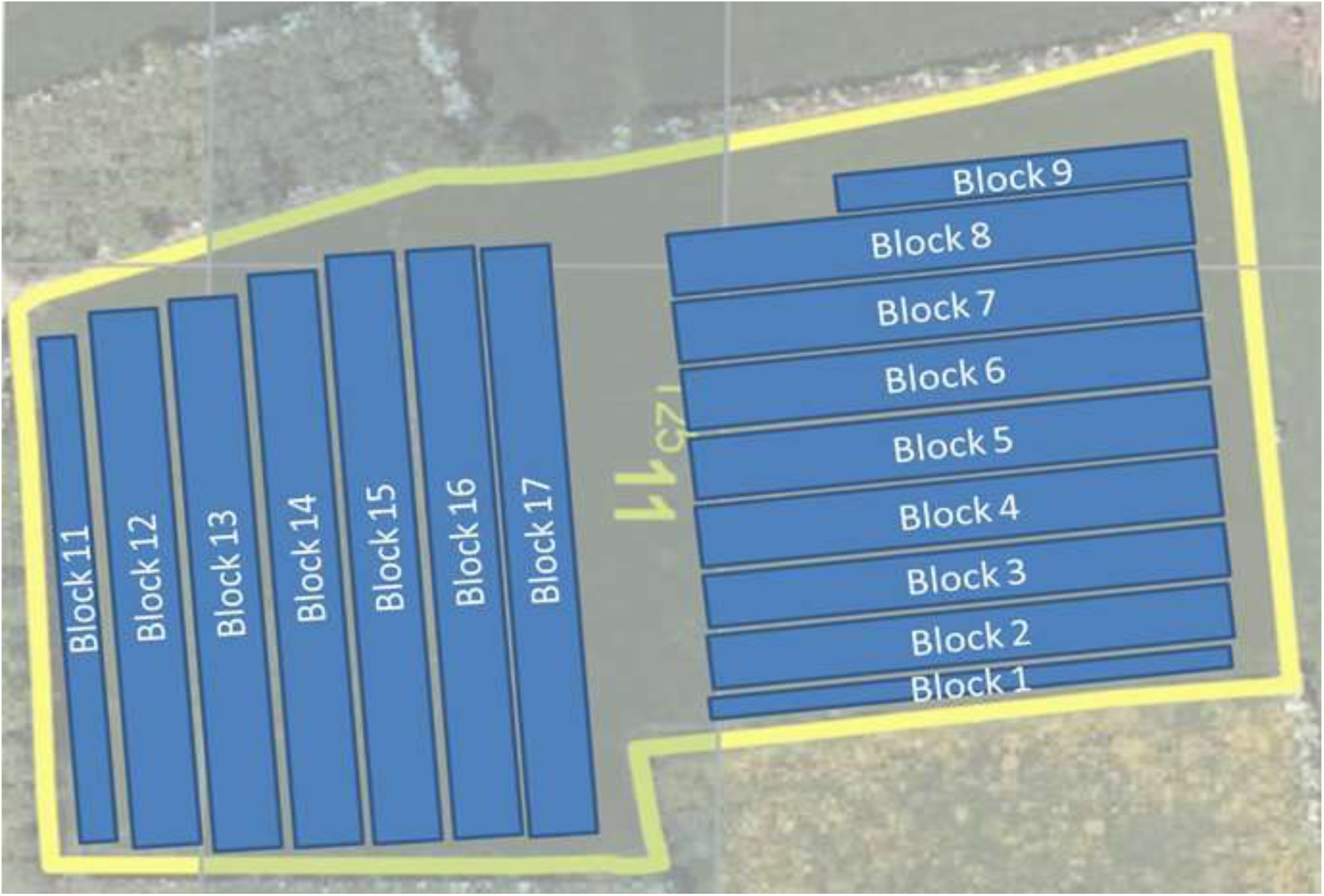
Location of field crop AFBI Loughgall breeding unit and Farquhar’s field

### Continued testing post December 2016

Since December 2016, a total of 718 further samples have been tested for PSTVd. These include 142 samples from the wider potato industry in Northern Ireland, and 650 from glasshouse protected breeding material in AFBI Loughgall. All of these samples have tested negative for PSTVd.

## Discussion

After the initial report of the PSTVd infected stock originating from AFBI Loughgall (EPPO RS 2016), testing of over 590 samples revealed that PSTVd was present in potato tubers in a field crop and in true potato seed in storage in AFBI Loughgall. An eradication programme following the original outbreak, followed by extensive further testing of 718 samples has not identified PSTVd infection in potato in Northern Ireland. Although the exact source of the PSTV infection in the AFBI Loughgall collection could not be conclusively identified, this research suggests that salt-tolerant breeding lines imported in the early 1980s are the likely source of the original infection. The presence of PSTVd in true seed in potato germplasm collections was first suggested by Camack and Harris (1973, EPPO Bulletin), following the first detection of PSTVd in the UK in true seed of potato in the early 1970’s (SPBS 1971; EPPO RS 1974). At the time the authors concluded that the presence of PSTV in breeding lines at this centre was probably not unique, and that many other collections that contained breeding lines possibly had low levels of infected material (Camack and Harris 1973). Further infections of PSTVd in potato germplasm were identified in Scotland by Harris and James (1987). Even up until 1993, many of the potato germplasm collections had not been fully tested for PSTVd by early 1993 (Jeffries et al. 1993). A 2014 outbreak of PSTVd in the Netherlands has also been traced back to infected potato germplasm (Verhoeven et al. 2018). Several others have noted the risk posed to global potato health by pathogen infected potato germplasm collections (Howell 1981; Jeffries et al. 1993; Owens and Verhoeven. 2009). In the case of the salt-tolerant material linked with the pathogen (DT02 and DT028) in the Northern Ireland detection, it is possible that the pathogen was at too low a level to be detected by the methods previously used to detect PSTVd in potato germplasm. Difficulties in detecting ‘mild strains’ using previously recommended methods have been noted (EPPO RS 1974). Modifications to existing tests, and the addition of more sensitive modern tests (q-PCR) have been suggested for use in the newer protocols (EPPO 2004).

In this research there were several instances of putative transmission of PSTV from the infected parent (DT02, DT028) to its progeny (e.g. J3911/5 the infected line notified by the Netherlands). PSTVd is known to be transmitted via infected pollen or ovules (Grasmick & Slack, 1986; Singh et al., 1992), which can result in up to 100% of the resulting seed infected (Fernow et al., 1970; Singh, 1970). In Tomato, over 60% of seed from infected plants can carry the viroid (Simmons et al 2015). Surface contamination was recorded in 12 samples of true seed in this study, and the most likely explanation for this is that the mechanical equipment used to extract seed (blending of the berries to release the seed) came into contact with the infected seed and passed this residual contamination to other seeds. .PSTVd has been shown to remain infective even after several hours on hard surfaces (Mackie et al. 2015), and this was taken into account in the testing scheme in this research which sought to identify if cross-contamination occurred between true seed in the crossing house at the same time as infected seed. An instance of putative transmission of the pathogen between unrelated breeding lines (i.e. from DT028 to Dundrum × L2678/96 progeny) was identified in the crossing house in 1992. We also identified 12 instances of surface contamination of the true seed with PSTVd.

Given that PSTVd was possibly present in breeding material in the AFBI Loughgall for many years, its sudden emergence is worth commenting upon. Any material bred through AFBI Loughgall, which was close to commercialisation, would have been submitted to the potato quarantine unit at Science and Advice for Scottish Agriculture (SASA). At SASA the material was tested for PSTVd and cultured to generate nuclear stock. None of the material sent to SASA from AFBI Loughgall was ever found to be infected with PSTVd. Furthermore, no PSTVd positive plants have ever been found in extant material. It should also be noted that as a result of an unrelated outbreak of PSTVd in the Netherlands in 2014 (EPPO RS 2014/88), all AFBI material sent in 2014 to a breeder in the Netherlands was tested and found to be PSTVd free. It is interesting to consider the reason for the delayed emergence of PSTVd from the AFBI Loughgall material, despite it being putatively present in material at the location for over 30 years. One explanation is that potatoes exhibiting symptoms of the disease would have been disposed of as part of the selection processes that are carried out during potato breeding. Misshapen or deformed tubers would have been disposed of, thus removing symptomatic material (i.e. possibly infected) from the collection. This removal of symptomatic material would have occurred in all cases, except in the exceptional circumstance where a seedling is kept despite exhibiting unhealthy symptoms, as is the case where a specific trait (salt tolerance) is of prime consideration. Thus, the recent emergence of PSTVd from AFBI Loughgall is likely to be related to the retention of non-commercial breeding material.

Extensive surveys and tests on material from the field crop in this study indicated five infected breeding lines all in the same plot, involving two closely related breeding lines (i.e. most likely infected by their parent CTO28) and three infected breeding lines the result of possible in glasshouse and/or in-field transmission. PSTVd is known to be transmitted between plants and plant material via contamination of machinery or between plants in close contact (Verhoeven et al 2010; Manzer and Merriam 1961). Transmission via aphids can also occur in the presence of potato leaf roll virus (Syller et al. 1997). Phytosanitary procedures for the infested location have been applied according to the EPPO standard for PSTVd on potato (EPPO 2011 PM9/13). All plant material from the infected plots and plots adjacent to infected plots (i.e. plots 1, 2, 3, 4) was removed from the field and destroyed by deep burial (>2 meters below surface level; Ebbels 2003) under supervision of the plant health regulator for Northern Ireland (i.e. DAERA). No commercial potato production was carried out within a 4 km radius of the infested field. Tuber and/or leaf testing of the remaining plots in the area yielded no PSTVd positive samples, however as a precaution all known hosts of PSTVd in the area around the site of the infected plants (approximately 8,000 m^2^) were also destroyed and the field ploughed and planted with grass. Rogue potatoes that re-sprout in the field are being treated with herbicide.

This work has described the detection of PSTVd in Northern Ireland for the first time. The NPPO for Northern Ireland in cooperation with AFBI has instigated a rescue plan testing scheme to ensure that selected breeding lines from AFBI Loughgall are safe for release to customers. The testing scheme was informed by the EPPO standard method for post entry quarantine for potato (EPPO 2004 PM3/21). Briefly, advanced breeding material and other material from the field crop for which there is no seed available from other sources unrelated to the Northern Ireland outbreak will be tested twice for PSTVd at two different growth stages. Tubers will be cored and some of the cores will be tested using q-PCR. Next, the remaining cores will be planted in a glasshouse and grown on at 18°C until they reach 25cm at which point their leaves will be tested for PSTVd. In addition, many of these breeding lines had been tested previously as part of the testing scheme to delineate the outbreak area in the infected field. Therefore, all plants will be tested at least twice for PSTVd. Since December 2016, over 650 samples from AFBI Loughgall have been tested for PSTVd, all of which have been PSTVd free. Stringent biosecurity protocols have also been implemented in the AFBI Loughgall site, to reduce the risk of the pathogen being spread. Tests are also being carried out on any at-risk material to be exported. This has resulted in the testing of 76 samples since December 2016, of which all were PSTVd free. Given the extensive scheme of testing, and exhaustive research and trace-back and trace-forward analysis carried out following this detection, the current status of PSTVd in Northern Ireland has been reported as actionable, under eradication. It is planned that in the absence of further findings in tests up to 2018 (2 years after detection) and the continued absence of potato volunteer plants in the outbreak field in 2019, the status will be updated to eradicated along the lines of the case of the PSTVd detection in the Netherlands in 2014 (EPPO RS 2016).

## Acknowledgements

The authors are grateful to the staff of AFBI and DAERA for their assistance with this work. This work was funded under the DAERA – AFBI assigned work programme.

